# A Statistical Framework for Evolutionary Analysis of Recurrent Somatic Mutations in Cancers

**DOI:** 10.1101/2020.04.10.036095

**Authors:** Xun Gu

## Abstract

Current cancer genomics databases have accumulated millions of somatic mutations that remain to be further explored, faciltating enormous high throuput analyses to explore the underlying mechanisms that may contribute to malignant initiation or progression. In the context of over-dominant passenger mutations (unrelated to cancers), the challenge is to identify somatic mutations that are cancer-driving. Under the notion that carcinogenesis is a form of somatic-cell evolution, we developed a two-component mixture model that enables to accomplish the following analyses. (*i*) We formulated a quasi-likelihood approach to test whether the two-component model is significantly better than a single-component model, which can be used for new cancer gene predicting. (*ii*) We implemented an empirical Bayesian method to calculate the posterior probabilities of a site to be cancer-driving for all sites of a gene, which can be used for new driving site predicting. (*iii*) We developed a computational procedure to calculate the somatic selection intensity at driver sites and passenger sites, respectively, as well as site-specific profiles for all sites. Using these newly-developed methods, we comprehensively analyzed 294 known cancer genes based on The Cancer Genome Atlas (TCGA) database.

## Introduction

Cancer occurs in almost all metazoans in which mutated somatic cells proliferate without any control (Stratton et al. 2009; Vogelstein et al. 2013; Martincorena and Campbell 2015). Progression to cancer involves sustained proliferation, altered energy metabolism, and abnormal responses to signals controlling cell growth, adhesion, and differentiation (Hanahan and Weinberg 2011). Thanks to the advances of next-generation sequencing (NGS) technologies in cancer genomics, The Cancer Genome Atlas (TCGA) has accumulated millions cancer somatic mutations from nearly ten thousand tumor-normal pairs. With this massive amounts of cancer genomic data, researchers have the opportunity to gain somatic mutation landscapes of more than 30 tumor types in order to better understand cancer biology and improve cancer diagnosis and therapy (Bailey et al. 2018)

A number of computational tools are developed for predicting cancer driver genes that may contribute to malignant initiation or progression, which can be categorized into six main types: (*i*) mutation frequency based, such as MuSiC (Dees et al. 2012), MutSig2CV (Lawrence et al. 2013), (*ii*) feature based (functional impact, for instance), such as OncodriveFML (Mularoni et al. 2016) and CompositeDriver (Bailey et al. 2018), (*iii*) structure (domain) based, such as e-Driver (Porta-Pardo and Godzik 2014) and ActiveDriver (Reimand and Bader 2013), (*iv*) evolutionary selective pressure based, such as dNdScv (Martincorena et al. 2017) and CBaSE (Weghorn and Sunyaev 2017), *C_N_/C_S_* (Zou et al. 2017), (*v*) network or pathway based (such as DriverNet (Bashashati et al. 2012), HotNet2 (Reyna et al. 2018) and (*vi*) multi-omics based (such as OncoIMPACT (Bertrand et al. 2015), OncodriverROLE (Schroeder et al. 2014). With the help of these tools, a list of cancer driver genes has been generated. For instance, over 1,000 genes have been identified as potential drivers to carcinogenesis, based on the data collected from the Catalogue of Somatic Mutations in Cancer (COSMIC) database (Tate et al. 2019).

However, not all mutations occurring in cancer driver genes are driver mutations (Futreal et al. 2004; Kandoth et al. 2013; Lawrence et al. 2014). Each mutation in a cancer genome is classified as a driver or passenger mutation according to its contribution to cancer development. Driver mutations are mutations that confer a growth advantage to the cells and have been positively selected for during the evolution of cancer, whereas passenger mutations do not have these features. Due to the overwhelming number of passenger somatic mutations that are unrelated to cancer, a great challenge is how to identify genetic and/or epigenetic events with functional implications to carcinogenesis (Chang et al. 2016; Ng et al. 2018; Tokheim et al. 2019). There are many computational tools, such as SIFT (Ng and Henikoff 2002), CHASM (Carter et. al 2009), HotSpot3D (Niu et al. 2016), which are designed to predict the cancer-driving sites.

As hallmark features of cancers stem from the disruption of basic, conserved cellular regulatory mechanisms evolved during the metazoan evolution toward multicellularity, understanding of common mechanisms driving tumorigenesis may provide insights into the evolutionary origins of multicellularity (Weinberg 1983; Nesse and Williams 1996; Merlo *et al* 2006). On the other hand, the notion that carcinogenesis is a form of evolution at the level of somatic cells suggests that our understanding of cancer initiation and progression can benefit from molecular evolutionary approaches. Although the detail remains highly controversial, it has been anticipated that cross-talks between biological processes of different evolutionary origins may present applicable treatment strategies for cancer. However, our understanding of cancer-multicellularity is still at the infant stage, owing to the lack of vigorous methodology.

## Results

### Data availability

We extracted the cancer somatic mutations of 10,224 cancer donors from TCGA PanCanAtlas MC3 project (https://gdc.cancer.gov/about-data/publications/mc3-2017), which includes 3,517,790 somatic mutations in the coding regions, 2,035,693 of which causing amino acid changes in cancers (missense mutations). After filtering out some redundancies according to the criterion of Bailey *et al*. (2018), this left us with 9,078 samples with 1,461,387 somatic mutations in the coding regions, which consisted of 793,577 missense mutations that were used in our study (Supplementary Table S1). Sequences and annotations of human genes were extracted from the Ensembl database (http://www.ensembl.org) (Aken et al. 2017). For each gene, we chose the transcript which is most commonly used in TCGA.

A list of 538 known cancer genes (Supplementary Table S2) was compiled by merging a list of 253 cancer genes with the missense mutations from the Cancer Gene Census Tier 1 (COSMIC GRCh37, V89) (Forbes et al. 2017; Tate et al. 2018), 127 significant mutated genes (SMGs) reported by Kandoth *et al*. (2013), 260 SMGs reported by Lawrence *et al*. (2014) and 299 cancer driver genes reported by Bailey *et al*. (2018).

### Two-component mixture model to distinguish drivers and passengers

To conduct a large-scale evolutionary analysis of cancer somatic mutations, we have to statistically distinguish between cancer-driving (amino acid) sites and passengers. It is known that passenger mutations may occur at amino acid sites of a gene randomly, with a very low recurrent frequency. Consequently, passenger sites, i.e., amino acid sites that are not cancer-driving, are highly unlikely to have recurrent somatic mutations at the same site across cancer samples. By contrast, cancer-driving mutations occur recurrently at only a few sites (called driver sires) across cancer samples (Stratton et al. 2009; Vogelstein et al. 2013; Martincorena and Campbell 2015).

We formulated a general framework of the two-component mixture model to distinguish between cancer driver sites and passenger sites of a gene, under which each site is either driver or passenger. The probability of any site being a driver is *η*, whereas that of being a passenger is 1-*η*. We denote *z* to be the number of somatic missense mutations (causing amino acid replacements) at an amino acid site. Let *P_0_(z)* be the distribution of *z* at a passenger site, and *P_1_(z)* is that at a driver site, respectively. It follows that the distribution of *z* of a gene under study is given by

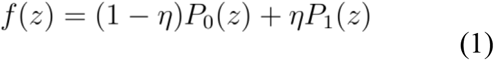

As somatic mutations at passenger sites are neutral or harmful for cancer progression (Ng and Henikoff 2002, Carter et. al 2009, Ng et al. 2018), it is biologically reasonable to assume a Poisson distribution for *P_0_(z)*, that is, 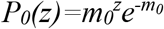, where *m_0_* is the recurrent rate at a passenger site. On the other hand, cancer somatic mutations occurred at driver sites are highly recurrent, subject to positive selection pressures in cancer progression that may differ considerably among sites (Futreal et al. 2004; Kandoth et al. 2013; McFarland et al. 2013; Lawrence et al. 2014). Therefore, a simple Poisson model may not be sufficient to account for the complexity of driver mutations. We thus develop a more realistic model as follows. First, at a given driver site, the number of recurrent mutations follow a Poisson process. Second, the recurrent rate at the driver site is modeled by *m_0_*+λ: while *m_0_* is the recurrent rate at a passenger site, λ is a random variable that varies among different driver sites according to a gamma distribution φ(λ)

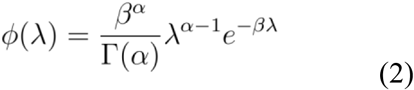

where α is the shape parameter and β is a scalar, and the mean recurrent rate at driver sites is *m_1_*=*m_0_*+α/β. It follows that the distributions of *z* at a driver site is given by

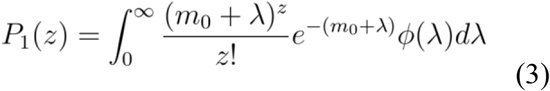

Note that *P_1_(z)* is reduced to a Poisson model when α→∞. Though the exact analytical form of *P_1_(z)* in Eq.(3), called the generalized negative binomial distribution (NBD*) is mathematically tedious, one can show that *P_1_(z)* approximately follows a negative binomial distribution (NBD) when *m_1_*>>*m_0_*, that is,

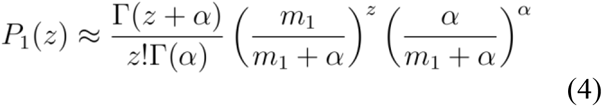

### Estimation procedure for the Poisson-NBD* model

We have developed a computational pipeline to estimate model parameters in the Poisson-NBD* model, as illustrated by the tumor suppresser gene PTEN. The cancer somatic mutation data were form TCGA database: among 403 amino acid sites of *PTEN*, 122 sites have at least one somatic missense mutation, and 59 of 122 sites have only one mutation **(Fig. 1A)**.

**Fig. 1.**
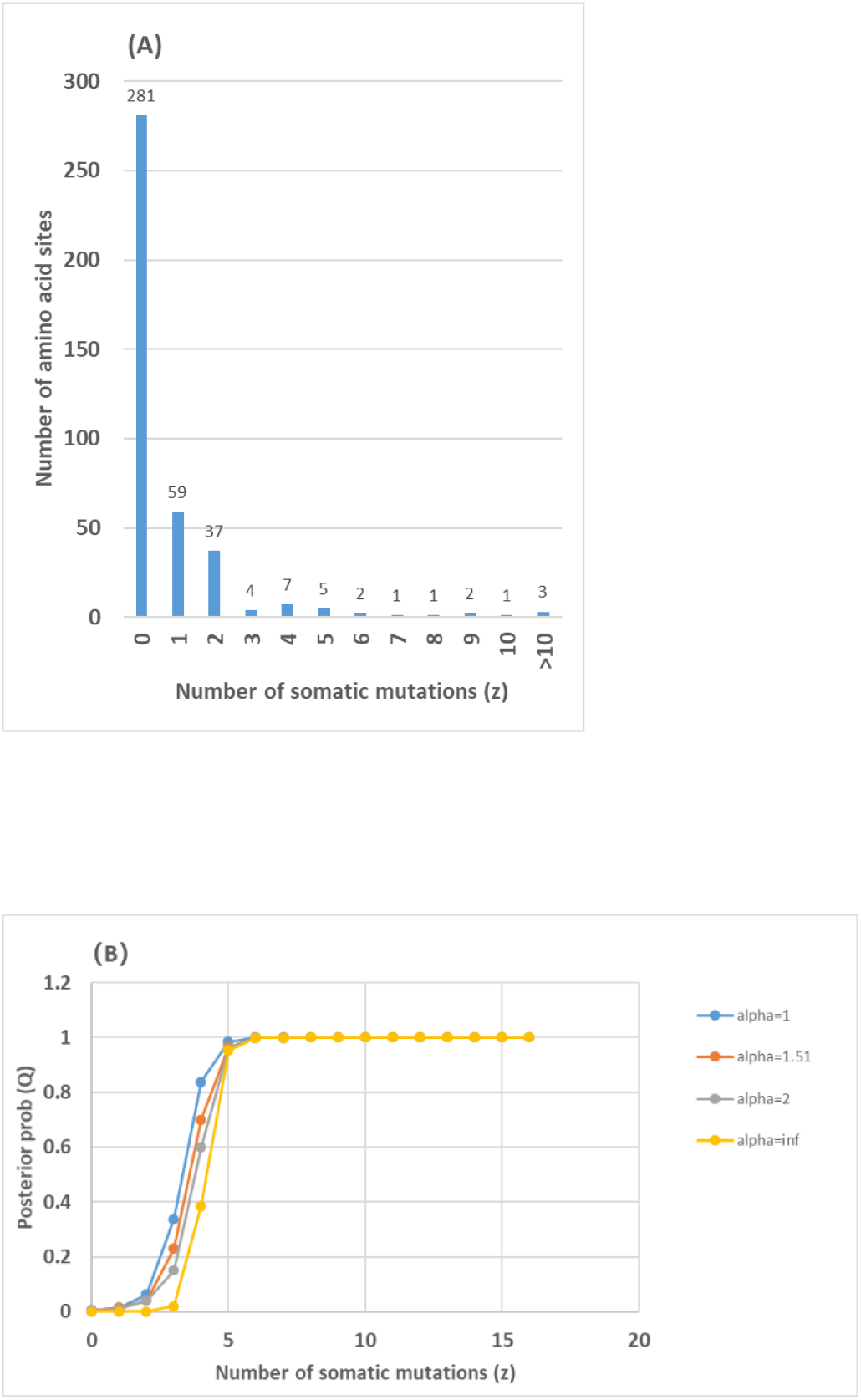
Analysis of tumor suppressor gene PTEN. (A) The histogram of the number of amino acid sites plotting against the counts of somatic missense mutations (*z*); one site for *z*=10, 12 and 60, respectively. (B) The posterior probability for the site to be cancer-deriving plotting against the counts of somatic missense mutations (*z*) under various values of the shape parameter: α=1, 1.51, 2 and ∞, respectively.

#### (i) Checking-point for single Poisson model as prerequisite

Count the number (*z*) of somatic mutations of each amino acid site of a gene, and then calculate the sampling mean 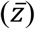 and variance [*var*(*z*)] of *z*, respectively. If *var*(*z*) is less than or equal to 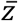, the algorithm will return to the output, claiming that a single Poisson model is sufficient to explain the pattern of somatic mutations. Further analysis is based on the condition of 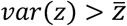. In the case of PTEN, we calculated 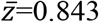 and *var*(*z*)=11.47, which apparently satisfies this criterion.

#### (ii) Estimation of m_0_ under the rare-driver assumption

For a given gene, let *f_0_* be the observed proportion of sites with no somatic mutation. By equating *f*(*z*) of Eq.(1) at *z*=0 with *f_0_*, one can write

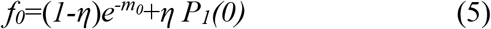

which can be further approximated by 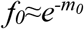 under the assumptions of *η*<<0.05 (very few cancer-driving sites) and *P_1_*(0)→*0* (sites without observed somatic mutations highly unlikely to be cancer-driving s). Thus, the first-order approximation to estimate *m_0_* is given by

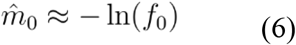

In the case of PTEN, we have *f0*=0.697 so that *m_0_* is estimated to be 0.361.

#### (iii) Determination of the shape parameter α

As shown by PTEN (Fig.1A), the empirical frequency of somatic mutations occurred in a single gene is usually highly skewed. Consequently, it is statistically difficult to estimate the shape parameter (α) in the NBD model of driver sites. To overcome this difficulty, we assume that the shaper parameter α is a universal parameter for most cancer genes so that one can estimate this parameter by the pooled cancer somatic mutations. After fitting the pooled distribution from 294 cancer genes to the Poisson-NBD* model by the maximum likelihood (ML) approach, we obtained the ML estimate α=1.51. In practice, one may also use a pre-specified α for model reduction: for instance, α=1 for the Poisson-geometric model, or α=∞ for the Poisson-Poisson model.

#### (iii) Simple method for estimating η and m_1_

One can easily derive the mean and variance of *z* under the Poisson-NBD* mixture model. By equating them with sampling mean and variance, respectively, we obtain the following relationships

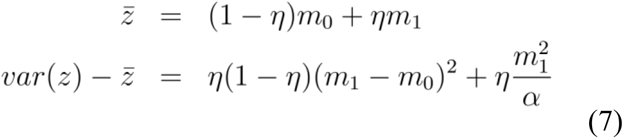

As long as the shape parameter α is given, we show that the MM (method of moments) estimate of *m_1_* is given by

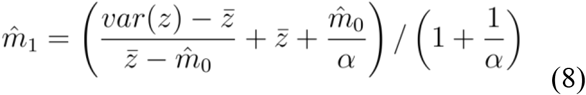

as well as the MM estimate of *η* is given by

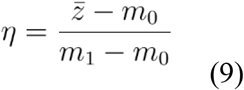

A nice property of Eq.(8) and (9) is that when the condition 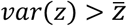 holds, the estimate of *η* satisfies 0<*η*<1, and the estimate of *m_1_* satisfies 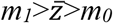. For PTEN, we estimated *m_1_*=13.89 (given α=1.51). The range of estimated *m_1_* is from 11.60 (α=1.0) to 22.9 (α=∞), suggesting a much higher rate of recurrent somatic mutations at driver sites than at passengers (*m_0_*=0.361). Meanwhile, the proportion of driver sites in PTEN was estimated as *η*=0.0356 (α= 1.51), with a range from 0.0215 (α=∞) to 0.0429 (α= 1.0). Hence, the number of driver sites of *PTEN* is expected to be 403×0.0356≈14.3, with a range from 8.7 (α=∞) to 17.3 (α=1.0), roughly consistent with the experimentally confirmed cancer-driving sites.

#### (iv) The quasi-likelihood test for η>0

Under the two-component mixture model, it is fundamental to test whether the proportion (*η*) of driver sites is significantly larger than 0. From the integrated cancer genomics databases, one can count the number of somatic mutations (*z_k_*) at each site *k* of a gene. The likelihood function based on Eq.(1) can be conventionally formulated as *L*=П_*k*_*f*(*z_k_*). Because of the highly skewed feature of cancer somatic mutations of a gene, we implement a simple ‘quasi-likelihood’ approach as follows. First, treat all model parameters except for *η* as known, either estimated or specified. Second, after two distributions *P_0_*(*z*) and *P_1_*(*z*) are then specified, one can easily show that the ML estimate of *η* based on Eq.(1) satisfies the following likelihood equation

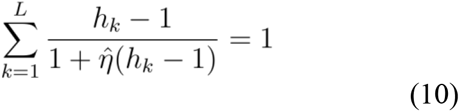

where *h_k_*=*P_1_*(*z_k_*)/*P_0_*(*z_k_*). Under the Poisson-NBD* mixture model, it is given by

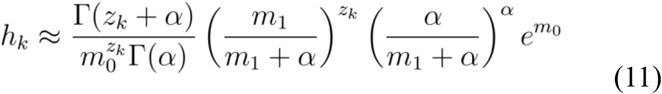

Two special cases needs to mentioned: when α=∞, i.e., the Poisson-Poisson mixture model, Eq.(11) is simplified to be

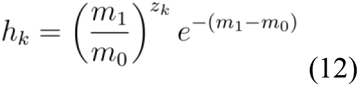

and when α=1, i.e., the Poisson-geometric mixture model, Eq.(11) is simplified to be

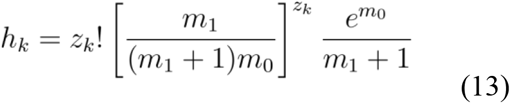

Third, an iterative algorithm is developed to obtain the ML estimate of *η* by numerically solving Eq.(10), with parameters *m_0_, m_1_* and α replaced by their estimates or specified values, and the MM estimate of *η* given by Eq.(9) is used as the initial value. And finally, an approximate likelihood ratio test (LRT) is designed to assert whether a two-component model is significantly better than a single-component model (*η*=0). The calculated log-LRT value approximately follows a chi-square distribution. Our computer simulation shows that the degree of freedom *df*=1 tends to be liberal, whereas *df*=2 tender to be conservative. In this sense, we recommend *df*=2 in practice. For *PTEN*, the null hypothesis of *η*=0 was statistically highly rejected (LRT test, *p*<10^-10^).

### Calculation of cellular selection for driver and passenger sites

The *C_N_/C_S_* ratio of a gene, or *Ω* for short, is defined by the ratio of the nonsynonymous mutation rate to the synonymous mutation rate in cancer samples. This ratio has been widely used in cancer genomic analyses as an indicator for the role of cellular-specific positive selection of an encoding gene in carcinogenesis (Bailey et al. 2018). A number of computation method were proposed to calculate the *C_N_/C_S_* ratio of a gene (Martincorena et al. 2017; and CBaSE (Weghorn and Sunyaev 2017), *C_N_/C_S_* (Zou et al. 2017). However, these methods did not distinguish between cancer-driving sites and passenger sites; a problem can be solved under the two-component mixture model. Let *Ω_0_* or *Ω_1_* be the *C_N_/C_S_* ratio of passenger sites or driver sites of a gene, respectively. We developed a simple method to calculate *Ω_0_* and *Ω_1_*, based on two assumptions: (*i*) *Ω_0_* is proportional to *m_0_*, the recurrent rate of somatic mutations at a passenger site, while *Ω_1_* is proportional to *m_1_*, the recurrent rate of somatic mutations at a driver site, and (*ii*) given the *C_N_/C_S_* ratio calculated from all sites of a gene, the constraint

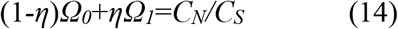

holds. After some basic algebras, we have the estimates of *Ω_0_* and *Ω_1_* as follows

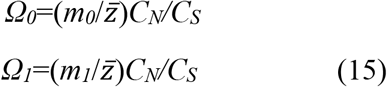

For instance, *Ω_0_*=4.77 and *Ω_1_*=183.5 in the case of PTEN. Moreover, one can verify that these estimates of *Ω_0_* and *Ω_1_* satisfy the constraint of Eq.(14) because 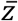, the mean number of somatic mutations per site, is expected to be 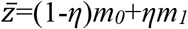.

### Statistical analysis of cancer genes

For 538 known cancer genes we compiled, 294 cancer genes are suitable for the two-component mixture model; the rest of genes have too little somatic mutations to be statistically reliable. Table 1 shows the statistical summary of three estimates (*m_0_*, *m_1_* and *η*) under various shape parameter values. First, the estimated proportion (*η*) of driver sites of a gene is usually very small, in a range from 0.4% (α=∞) to 1.0% (α=1). Second, while only less than 1% of sites of cancer genes are possibly cancer-driving, LTR tests showed the majority *η* estimates are significantly greater than 0, and the result is insensitive to the shaper parameter (α). Third, the number of somatic mutations per driver site (*m_1_*) is at least one magnitude larger than that per passenger site (*m_0_*), indicating much more recurrent somatic mutations occurred at driver sites than at passenger sites. In short, our analysis supports the notion that among numerous somatic mutations appearing in cancer samples, only a very small portion actually drives the process of carcinogenesis, subject to a very strong positive selection. Indeed, Table 1 shows that *Ω_1_* (the *C_N_/C_S_* ratio at driver sites) is at least one-magnitude higher than *Ω_0_* (the *C_N_/C_S_* ratio at passenger sites). For example, Table 2 shows the results of some well-known cancer genes.

**Table 1.** Summary for the statistical analysis of 294 cancer-related genes based on two-component (Poisson-NBD*) mixture model

| | $\alpha=1.0$ | $\alpha=1.51$ | $\alpha=2$ | $\alpha=\infty$ |
| --- | --- | --- | --- | --- |
| $\eta$ | (0.987±0.103)% | (0.820±0.022)% | (0.740±0.078)% | (0.396±0.038)% |
| $m_0$ | 0.095±0.003 | 0.095±0.003 | 0.095±0.003 | 0.095±0.003 |
| $m_1$ | 2.883±0.219 | 3.450±0.264 | 3.813±0.292 | 5.671±0.438 |
| $\Omega_0$ | 1.343±0.146 | 1.343±0.146 | 1.343±0.146 | 1.343±0.146 |
| $\Omega_1$ | 57.81±12.47 | 69.18±15.02 | 76.46±16.52 | 113.7±24.69 |
| *LRT test against the null hypothesis of $\eta=0$: number of genes* |  |  |  |  |
| $P<0.0001$ | 137 (47.2%) | 137 (47.2%) | 137 (47.2%) | 137 (47.2%) |
| $P<0.01$ | 173 (58.8%) | 170 (57.8%) | 170 (57.8%) | 169 (57.5%) |
| $P<0.05$ | 235 (79.9%) | 231 (78.6%) | 229 (77.9%) | 226 (76.9%) |

Note: Three parameters ( $m_0$ , $m_1$ and $\eta$ ) and two $C_N/C_S$ measures ( $\Omega_0$ and $\Omega_1$ ) were estimated under different values of the shape parameter $\alpha$ ; in this case the mean over 294 cancer genes were presented, follows by standard errors. For the LRT test against the null hypothesis of $\eta=0$ , the number of genes were presented under a given by p-value cutoff.

**Table 2.** Examples of some well-known cancer genes analyzed by the Poisson-NBD* mixture model.

| Gene | $m_0$ | $m_1$ | $\eta$ | $C_N/C_S$ | $\Omega_0$ | $\Omega_1$ |
| --- | --- | --- | --- | --- | --- | --- |
| NRAS | 0.106 | 11.343 | 0.0123 | 86.94 | 37.65 | 4030.6 |
| HRAS | 0.094 | 21.753 | 0.0257 | 27.65 | 4.00 | 924.3 |
| IDH1 | 0.080 | 2.633 | 0.0007 | 22.69 | 22.17 | 725.8 |
| KRAS | 0.148 | 25.788 | 0.0243 | 14.46 | 2.77 | 483.3 |
| BRAF | 0.076 | 6.776 | 0.0111 | 9.71 | 4.90 | 437.7 |
| AKT1 | 0.051 | 24.938 | 0.0042 | 2.05 | 0.67 | 326.5 |
| PIK3CA | 0.127 | 12.757 | 0.0272 | 11.27 | 3.03 | 305.8 |
| MAPK1 | 0.063 | 11.264 | 0.0040 | 2.17 | 1.26 | 225.6 |
| GNA11 | 0.040 | 21.461 | 0.0052 | 1.48 | 0.39 | 211.8 |
| PTEN | 0.361 | 13.879 | 0.0357 | 11.15 | 4.77 | 183.5 |
| FGFR3 | 0.064 | 16.564 | 0.0044 | 1.49 | 0.70 | 181.1 |
| SMAD4 | 0.134 | 8.959 | 0.0113 | 4.61 | 2.63 | 176.6 |
| ERBB2 | 0.065 | 12.800 | 0.0054 | 1.53 | 0.74 | 146.3 |
| FGFR2 | 0.100 | 7.883 | 0.0069 | 2.19 | 1.43 | 112.9 |
| EGFR | 0.116 | 9.757 | 0.0106 | 1.84 | 0.98 | 82.2 |
| RAF1 | 0.043 | 4.015 | 0.0025 | 0.93 | 0.76 | 71.5 |
| TP53 | 0.481 | 21.457 | 0.2008 | 12.20 | 1.25 | 55.8 |
| MYC | 0.097 | 3.502 | 0.0071 | 1.17 | 0.94 | 33.8 |
| NOTCH1 | 0.077 | 1.811 | 0.0069 | 1.49 | 1.29 | 30.3 |

Note: Estimates of three parameters ( $m_0$ , $m_1$ and $\eta$ ) are based on the model with $\alpha=1.51$ . $C_N/C_S$ is estimated by the method of Zhou et al. (2018).

### Posterior probability for a site being cancer-driving

An important application of the nearly-developed model is to provide a statistically sound method for scoring cancer-driving sites. This task can be achieved by implementing an empirical Bayesian approach to calculating *P*(*driver*|*z*), the posterior probability of an amino acid site being a cancer-driver when the number (*z*) of somatic mutations is observed. For a given gene, we consider *π*(*driver*)=*η* to be the prior probability of an amino acid site being a driver. Let *Q_k_*=*P*(driver|*z_k_*) be the posterior probability of being a driver at the *k*-th site of a gene with *z_k_* observed somatic mutations. The Bayesian rule claims that

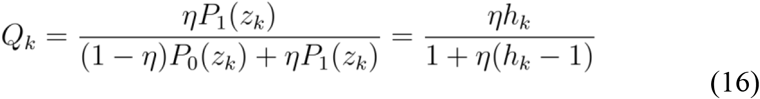

After three model parameters (α, *m_0_, m_1_* for calculating *h_k_*) as well as *η* are replaced by their estimates (or specified values), *Q_k_* can be calculated according to Eq.(16) by the means of empirical Bayes. The relationship between *Q_k_* and *z* is logistic; the baseline of *Q_k_* is determined by *z_k_*=0, which is usually close to zero. A higher *Q* value indicates a higher statistical confidence for cancer somatic mutation sites to be cancer-driving. As the number of observed somatic mutations (*z_k_*) at a site increases, the chance of this site to be a driver, measured by *Q_k_*, increases toward to 100% ultimately.

**Fig.1B** shows the posterior probability of being a driver site plotting against the number (*z*) of somatic mutation at a site of PTEN gene, under various shape parameter values, i.e., α=1, 1.51, 2 and ∞, respectively. Amino acid sites with at least five recurrent mutations (z⩾5) have high (>0.95) posterior probabilities to be driver sites, regardless of the value of α; whereas sites with a few somatic mutations (*z*⩽2) are highly unlikely. In this case, one should be cautious for those sites with *z*=3 or 4 because the posterior profiling is highly dependent of the shape parameter α. Since a larger α value tends to give a lower posterior probability, i.e., to be more conserved, the Poisson-Poisson model (α=∞) is suggested if a posterior probability cutoff of 0.5 is used to predict cancer-driving sites. Yet, we recommended a more stringent cutoff as high as 0.95 that is insensitive to the model selection.

### Site-specific posterior mean of somatic mutation rate

Because different driver sites may have different rates of recurrent somatic mutations in cancers, a single measure may not be sufficient to characterize the pattern of somatic mutations at driver sites. The Poisson-NBD* mixture model is flexible to implement a Bayesian approach to predicting site-specific posterior mean of the somatic mutation rate. Recall that the rate of recurrent mutation at a driver site is *m_0_*+λ, where λ is a random variable that varies among different driver sites according to a gamma distribution φ(λ) given by Eq.(2), with the mean α/β and the shape parameter α. Note that the mean of *P_1_*(z) is given by *m_1_*=*m_0_*+α/β. Hence, the posterior density of λ given *z* somatic mutations at a driver site is given by

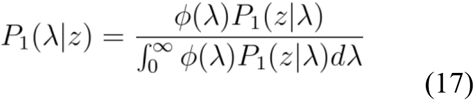

Though the exact result of *P_1_*(λ|*z*) is tedious, we derived a useful close form when *m_1_*>>*m_0_*; in this case *P_1_*(λ|*z*) approximately follows a gamma distribution, with the mean conditional of *z* is given by

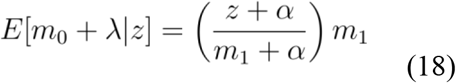

Thus, given *z_k_* somatic mutations at the *k*-th site of a gene, the posterior mean of recurrent mutations at site *k* is given by *Mk*=(1-*Q_k_*)*m_0_*+*Q_k_E*[*m_0_*+λ|*z_k_*]. Together with Eq.(18), we have

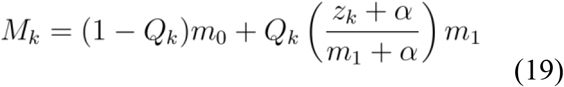

It should be noticed that the variability of site-specific *M_k_*-profile highly depends on the shaper parameter α. When α is small, the site-specific *M_k_*-profile varies dramatically with the number of somatic mutations (*z_k_*). By contrast, when α is very lager, it virtually becomes an average of *m_0_* and *m_1_* weighted by the (site-specific) posterior probability.

### Site-specific posterior C_N_/C_S_ profiling

Eq.(19) provides an elegant site-specific profile to demonstrate the pattern of somatic mutations in cancers. However, the magnitude of the variation of *M_k_* among sites highly depends on the sample size, making the follow-up analysis difficult when various datasets with different sample sizes are compared. It is therefore desirable to develop a site-specific *C_N_/C_S_* profile that may indicate the underlying selection mechanism on cancer somatic mutations. Let *Ω\** be the posterior mean of *C_N_/C_S_* ratio at a driver site, with *z* somatic mutations observed. In analogy to Eq.(18), one may write

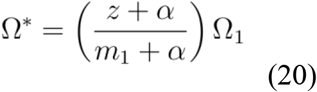

where *Ω_1_* is the mean *C_N_/C_S_* ratio over driver sites of a gene given by Eq.(15). Let *Ω_k_* be the posterior mean of *C_N_/C_S_* ratio at site *k*, with *z_k_* somatic mutations observed. Naturally one may write *Ω_k_*=[1-*Q*(*z_k_*)]*Ω_0_*+*Q*(*z_k_*)*Ω\**, where *Ω_0_* is given by Eq.(15). Together with Eq.(20), we have

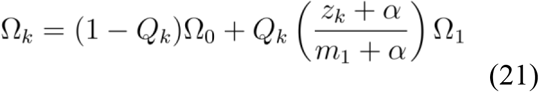

**Fig.2** shows the site-specific *Ω_k_*-profile of PTEN for α=1.51 (panel A) and α=∞ (panel B), respectively. Similar to the α-dependency of site-specific *M_k_*-profile, the site-specific *Ω_k_*-profile with α=1.51 reveals a high variability of selection pressures among sites, whereas that with α=∞ reveals virtually a binary-like pattern.

**Fig. 2.**
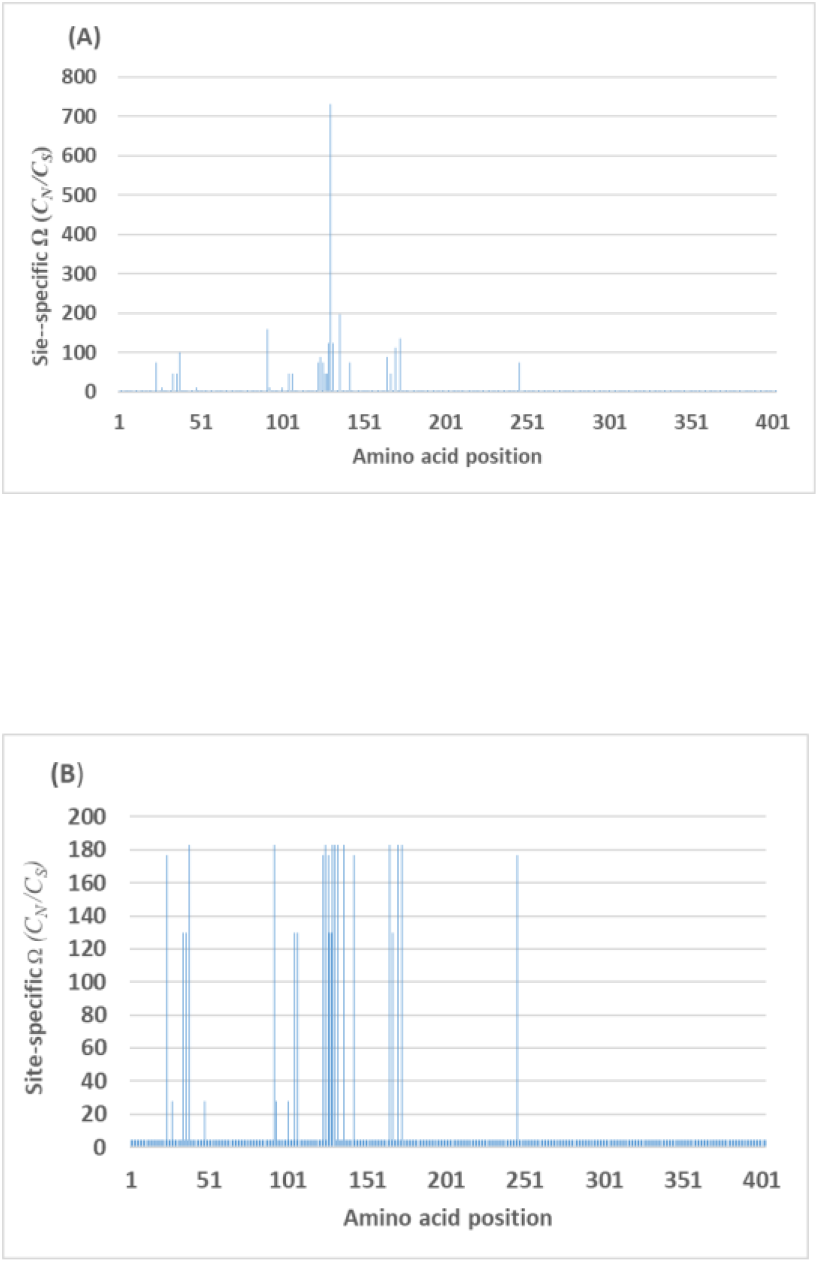
Site-specific *Ω_k_*-profile of PTEN plotting against the amino acid sequence, where α=1.51 (panel A) and α=∞ (panel B), respectively.

### Software availability

The methods we have reported above are integrated to the software (*CanDriS*, short for <u>Can</u>cer <u>Dri</u>ving <u>S</u>ite analysis) designed under the Perl environment, which is available on the GitHub (https://github.com/jiujiezz/candris).

## Discussion

### Prediction of putative cancer genes

The two-component mixture model can be used to predict cancer genes. The current version implemented a quasi-likelihood approach to estimate *η*, the proportion of driver sites of a gene. If the quasi-likelihood ratio test rejected the null hypothesis of *η=0*, one may predict a gene as a putative cancer-related gene, because the single component model is statistically not supported. Indeed, our data analysis showed that at least 75% known cancer genes showed *η>0* significantly (*p*<0.05, see Table 1). Yet, it remains unclear whether it is appropriate for making some new cancer-gene predictions, due to the small sampling bias as well as model simplicity. At any rate, one should be cautious for cancer-gene predicting without further experimental validations.

### Prediction of cancer-driving sites

In cancer genomics, the number (*z*) of recurrent somatic mutations at a given site is one of the primary indicators for the driving-role in carcinogenesis. While it is intuitive to predict a driver site when a large counts of recurrent somatic mutations have been observed, it is difficult to choose an objective cutoff because it is sample size-dependent, especially for those sites with an intermediate number of somatic mutations. The two-component mixture model we reported here avoids these drawbacks by calculating the posterior probability of a site being a driver, a statistically sound criterion for the driver site prediction.

Our analysis recovered almost well-known cancer-driver sites that feature a high frequency of recurrent mutations, including most driver sites in TS (tumor suppressor) genes with around 10-20 somatic mutations. An intriguing question is whether the posterior probability of a site being a driver can be used to predict amino acid sites that are weakly cancer-driving if the number (*z*) of recurrent mutations is intermediately low (*z*=5-10). As shown by Fig.1, calculation of the posterior probability in that range could be highly dependent of the model selection (measured by the shaper parameter α). Tentatively, we recommend that one may the Poisson-Poisson two component model, the most conserved method for predicting weak driver sites.

### Model selection and extension

Before the Poisson-NBD* model is applied for cancer somatic mutations of a gene, we have determine the shape parameter α. As this parameter cannot be reliably estimated from the distribution of somatic mutations of this gene, we have to specify α or estimate it based on a pooled dataset. The remaining question is to what extent such practice may affect the result of our analysis. As shown in Table 1, we found that the effects of α on key parameter estimates (*m_0_*, *m_1_* and *η*) generally do not alter their biological interpretations. Moreover, the posterior probability might be affected considerably when it is around 0.5 (Fig.1B), whereas it would be virtually no effect when the cutoff is set to be 0.75 or higher.

On the other hand, both site-specific profiles of *M_k_* and *Ω_k_* given by Eq.(19) and Eq.(21), respectively, are sensitive to the choose of α. When α=∞, *M_k_* is virtually the mean of *m_0_* and *m_1_* weighted by the posterior probability at site *k*, and *Ω_k_* the mean of *Ω_0_* and *Ω_1_*. As α decreases, the magnitudes of their variation among sites with *z_k_* dramatically, as illustrated in Fig.2. Since the estimate from the pooled cancer gene data showed α=1.51, we recommend that α=1~2 would be appropriate in these analyses.

The Poisson-NBD* model can be extended to include several biological covariates that may have nontrivial effects. Cancer samples include different cancer stages (primary or metastasis), clinical severity, type of cancers, and medical treatments. Further study will take these factors into account in the two-component model. For instance, one may improve our computational pipeline by implementing more options and flexibility for different types of cancer samples. Meanwhile, the pattern of somatic mutations are highly heterogeneous among chromosome regions (Lawrence et al. 2013, Kandon et al. 2013; Alexandrov et al. 2013); hence the Poisson model for passengers may be biologically oversimplified. With these caveats in mind, our study showed that the two-component Poisson-NBD* mixture model is useful for cancer genomics analysis.

### Technical treatment for some extremes

Cancer-related genes may have highly recurrent somatic mutations at very few sites (hotspots), e.g., the 600^th^ codon of *BRAF*, the highest recurrent mutation site, has received 545 recurrent mutations, much higher than the second site (469^th^ codon) with 16 mutations. In our dataset, there are six well-known cancer-related genes (including oncogenes *KRAS, BRAF, IDH1, PIK3CA, NRAS* and tumor suppressor gene *TP53*), each of which has at least one site with more than 100 recurrent somatic mutations. While those highly recurrent mutations are apparently cancer-related, they may cause some statistical biases in our prediction (not shown). Therefore, we excluded any amino acid site with *z*>100 in our analysis, and assigned the posterior probability of being a driver site *Q*(*z*>100)=1.

## Notes

### Competing Interest Statement

The authors have declared no competing interest.

